# Sleep homeostasis in lizards and the role of cortex

**DOI:** 10.1101/2024.07.31.605950

**Authors:** Sena Hatori, Sho T. Yamaguchi, Riho Kobayashi, Futaba Matsui, Zhiwen Zhou, Hiroaki Norimoto

## Abstract

Although their phenotypes are diverse, slow-wave sleep (SWS) and rapid eye movement sleep (REMS) are the two primary components of electrophysiological sleep (e-sleep) in mammals and birds. Slow waves in the cortex not only characterize SWS but are also used as biological markers for sleep homeostasis, given their rebound after sleep deprivation (SD). Recently, it has been reported that the Australian dragon *Pogona vitticeps* exhibits two-stage sleep pattern in the dorsal ventricular ridge (DVR), which includes a homologue of the mammalian claustrum (CLA). It remains unclear whether reptilian e-sleep, which has been characterized by activity outside the cortex, compensates for sleep loss, as observed in mammals. We here report a significant rebound in the local field potential (LFP) after 7 hours of SD, during both SWS and REMS. Meanwhile, the cycle and mean bout length of SWS/REMS remained unaffected. We further investigated a possible role of the cortex in e-sleep regulation and homeostasis in *Pogona* and found that, although a corticotomy had no obvious effect on the LFP features investigated, it abolished LFP power rebound in the CLA/DVR after SD. These findings suggest that e-sleep homeostasis is a common feature in amniotes, and that cortex is involved in regulating activity rebounds in reptiles and mammals.

## Introduction

Sleep is a fundamental physiological process observed in various animals and is regulated in a homeostatic manner. Slow-wave sleep (SWS) and rapid eye movement sleep (REMS) are the two primary components of electrophysiological sleep (e-sleep) in mammals and birds^1^. In mammals, the sleep state is determined by electroencephalogram (EEG) recordings, which are acquired from the cortex. It has been reported that sleep deprivation (SD) leads to a strong increase in the intensity of cortical delta (δ) -band (0.1-4 Hz) activity during subsequent SWS, thus making it a reliable marker of sleep need^2,3^. This elevated δ power decreases progressively with subsequent sleep. It has been reported that δ power activity facilitates neuroplastic changes, potentially playing a compensatory role for sleep loss^4^. Therefore, examining e-sleep homeostasis could provide insights into the function and significance of sleep.

Recent studies have revealed that reptiles including *Pogona vitticeps* have SWS and REMS, suggesting that these two sleep stages may have originated before the diversification of amniotes 320 million years ago^5,6^. *Pogona* takes a monophasic sleep, in which SWS and REMS are repeated within a 2-min cycle. These two modes are classified according to local field potential (LFP) from dorsal ventricular ridge (DVR), outside the cortex. The SWS in *Pogona* is characterized by the irregular occurrence of δ-band oscillations called sharp-waves (0.1-4 Hz, ShW), which originates in the anterior medial pole of DVR, likely a homologous to the mammalian claustrum (CLA), and propagates to the neighbouring DVR^7^. This characteristic provides a powerful system to address some key issues regarding sleep homeostasis: (i) Are there commonalities in e-sleep homeostasis across amniotes? (ii) Does the rebound of δ-band activity occur in animals in which e-sleep has been determined based on the activity outside the cortex? (iii) If such a rebound occurs, does cortex play a role? We addressed these questions by recording sleep activity in the CLA/DVR after 7 hours of sleep deprivation.

## Results

### SWS in lizards is under homeostatic regulation

First, we examined the effects of SD on SWS. LFP recordings were obtained from either the CLA or adjacent DVR (Fig. 1A). During monophasic sleep at night, SWS and REMS alternated regularly, as previously described^5^ (Fig. S1). The lizards were continuously deprived of sleep for 7 hours by gentle handling and visual distraction, whereas control animals were allowed to sleep normally (Figs. 1B, C, Supplementary Movie 1). Comparing the values of δ power in SWS during Pre-sleep (Pre, day 1) and recovery sleep (RS, day 2) revealed a significant increase in δ power in the early phase of RS, followed by a gradual decline (Fig. 1E, Pre vs. RS, \*\*\**P* = 2.11 × 10^-5^, two-way ANOVA). Similarly, the δ power ratio, defined as the ratio of the δ power during RS to the δ power during Pre, increased in the SD group compared to the non-sleep deprivation (NSD) group (Fig. 1F, NSD vs. SD, \*\*\**P* = 4.3 × 10^-17^, two-way ANOVA; NSD vs. SD, \*\*\**P* < 1.0 × 10^-7^, bootstrap test for means). The observed increase in δ power following SD resulted from an increase in the amplitude of ShW, rather than the frequency (Figs. 1G-I, ShW amplitude ratio, NSD vs. SD, \*\*\**P* = 4.8 × 10^-17^, two-way ANOVA; NSD vs. SD, \*\*\**P* < 1.0 × 10^-7^, bootstrap test for means; ShW frequency, Pre vs. SD, *P* = 0.099, two-way ANOVA). Animals typically woke up immediately after the beginning of the light period after SD. However, the δ power rebound continued despite resuming the light period (ZT 0-1). This resembles “sleep inertia”, a transitional state of grogginess immediately after awakening from sleep^8,9^ (Fig. 1J, NSD vs SD, \*\*\**P* = 2.6 × 10^-4^, bootstrap test for means).

**Figure 1.**
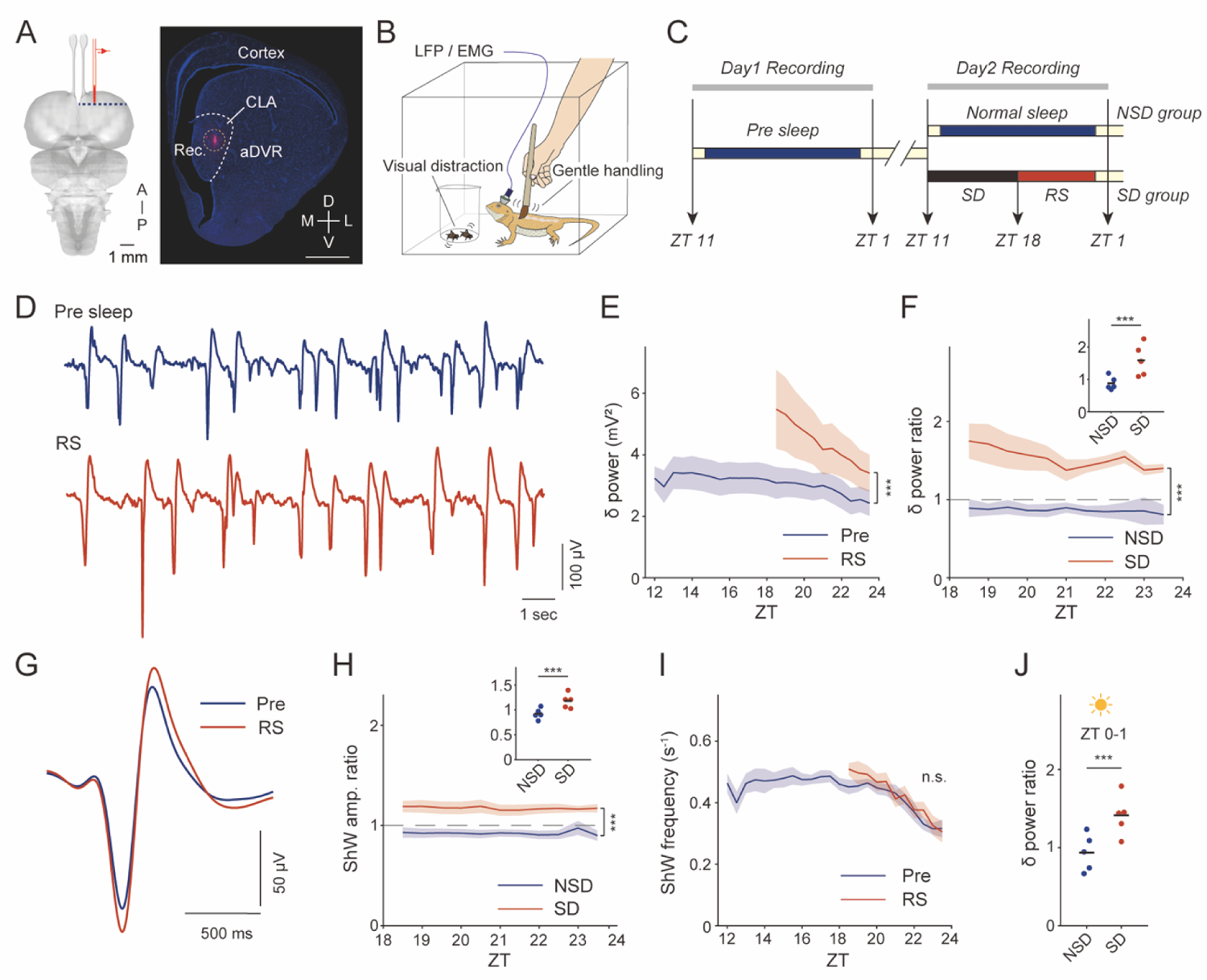
SWS in lizards is under homeostatic regulation. (A) Left: schematic diagram of the recording configuration. Coronal section outlined by a blue dotted line is shown on the right. Right: a corresponding coronal section with DAPI (blue) and DiI dye (red) to identify the electrode track. Scale bar, 1 mm. Rec, recording site; CLA, claustrum; aDVR, anterior dorsal ventricular ridge; A, anterior; P, posterior; D, dorsal; V, ventral; M, medial; L, lateral. (B) Schematic diagram of sleep deprivation experiment. Lizards underwent sleep deprivation (SD) by gentle handling and visual distraction for 7 hours. See also Supplementary Movie 1. (C) Time course of the experimental procedures. The non-sleep deprivation (NSD) group was designated as the control and normal sleep was recorded during two consecutive nights under standardized conditions. In the SD group, following Pre-sleep recording, the lizards underwent SD in ZT 11-18 on the subsequent day, and the succeeding recovery sleep (RS) was recorded. (D) Sample traces of Pre-sleep and RS during SWS. (E) δ power in SWS. N = 5 recordings from 5 animals each. Data are presented as mean ± SEM at 30 min intervals. (F) δ power ratio in SWS. δ power ratio is calculated by dividing δ power during RS by δ power during Pre-sleep. N = 5 sessions from 3 animals in NSD and 5 animals in SD. Data are presented as mean ± SEM at 30 min intervals. (Upper right) δ power ratio in ZT 18.5-21.5. N = 5 sessions from 3 animals in NSD and 5 animals in SD. (G) Averaged traces of sharp waves (ShWs) during SWS from one animal. All ShWs were detected from ZT 18.5-21.5. N = 2290 (Pre) and 1984 (SD) respectively. The mean ± SEM are plotted. (H) ShW amplitude ratio in SWS. The ratio is calculated by dividing the ShW amplitude during RS by the ShW amplitude during Pre-sleep. N = 5 sessions from 3 animals in NSD and 5 animals in SD. Data are presented as mean ± SEM at 30 min intervals. (Upper right) ShW amplitude ratio in ZT 18.5-21.5. N = 5 sessions from 3 animals in NSD and 5 animals in SD. (I) ShW frequency in SWS. N = 5 recordings each from 5 animals. The data are presented as mean ± SEM at 30 min intervals. (J) δ power ratio during the following morning (ZT 0-1). N = 5 sessions from 3 animals in NSD and 5 animals in SD.

### Synaptic efficacy is enhanced after SD

To investigate whether the changes in synaptic transmission occur in the CLA/DVR after SD, we performed whole-cell patch clamp recordings. We harvested coronal CLA/DVR slices (600 μm) from the lizards that had been sleep deprived for 7 hours (Fig. 2A). Miniature excitatory postsynaptic currents (mEPSCs) were recorded from the excitatory cells, and the frequency and amplitude of mEPSCs were analyzed. We found that the frequency and amplitude of mEPSCs are significantly higher in the lizards that underwent SD compared to the naturally sleeping lizards (Figs. 2C, D, mEPSC frequency, Control vs. SD, \**P* = 0.044, bootstrap test for means; mEPSC amplitude, Control vs. SD, \*\*\**P* < 1.0 × 10^-7^, *D* = 0.30, Kolmogorov-Smirnov test), indicating that sleep down-regulates the synaptic efficacy also in the CLA/DVR.

**Figure 2.**
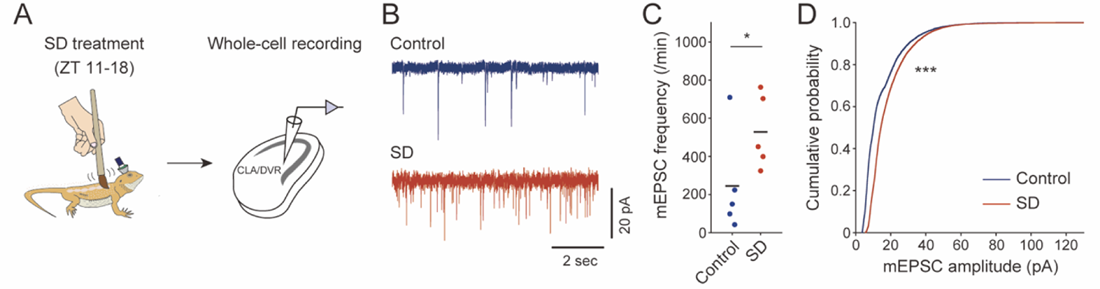
Synaptic efficacy of glutamatergic neurons is enhanced after SD. (A) Schematic illustration of experimental procedures. (B) Representative traces of mEPSCs. (C) Changes in the frequency of mEPSCs. N = 5 slices from 3 animals each. (D) Cumulative probability of mEPSCs amplitude. N = 9113 (Control) and 13120 (SD) mEPSCs.

### REMS in lizards is also homeostatically regulated

Next, we investigated LFP rebound during REMS. In mammals and birds, REMS homeostasis is reflected by the theta (4-12 Hz) power during REMS^10^. However, unlike SWS, there is no consensus regarding REMS homeostasis; the results vary depending on the species, or even within the same species, owing to different methodologies^2,11,12,13,14,15,16,17^. In *Pogona*, REMS is characterized by broadband energy measured in beta (β) power (10-30 Hz) in the CLA/DVR. Therefore, we examined the effect of SD on β power during subsequent REMS. In the SD group, β power rebound was observed during the early phase of RS, but was diminished during the later phase and the following morning (Figs. 3B, C, β power ratio at night, NSD vs. SD, *** *P* = 1.2 × 10^-6^, two-way ANOVA; NSD vs. SD, \**P* = 0.01, bootstrap test for means; β power ratio in the morning, NSD vs. SD, *P* = 0.56, bootstrap test for means). These data suggest that REMS in *Pogona* is under homeostatic regulation.

**Figure 3.**
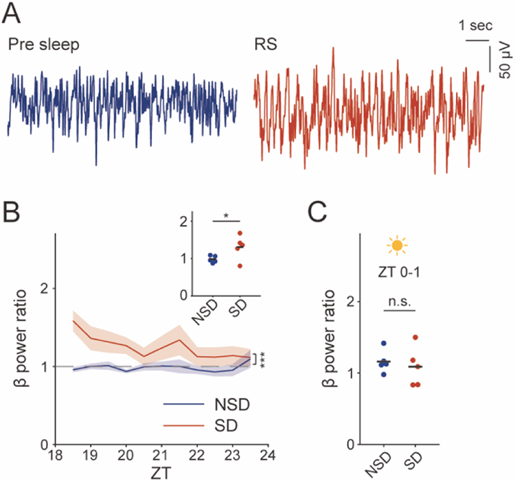
REMS in lizards is also homeostatically regulated. (A) Sample traces of Pre-sleep and RS during REMS. (B) β power ratio in REMS. N = 5 sessions from 3 animals in NSD and 5 animals in SD. Data are presented as mean ± SEM at 30 min intervals. Upper right: β power ratio in ZT 18.5-21.5. N = 5 sessions from 3 animals in NSD and 5 animals in SD. (C) β power ratio on the following morning (ZT 0-1). N = 5 sessions from 3 animals in NSD and 5 animals in SD.

### Sleep architecture is not affected by sleep deprivation

Diurnal animals exhibiting monophasic sleep tend to have species-specific cycles of SWS/REMS^5,6,18,19^. In *Pogona*, these two sleep modes alternate regularly within a 2 min-cycle approximately (Fig. 4A) and persist throughout the night. We investigated whether the regularity of SWS/REMS alternation was influenced by SD treatment. To assess the periodicity of these states, we used the autocorrelation function of the δ/β time series^5^, from which we extracted the oscillation period and peak-to-valley (P2V) difference, an index of the robustness of SWS/REMS alternation^6^ (Fig. 4B). During Pre, P2V levels remained elevated for approximately 8 hours during the dark period. The RS phase exhibited a peak in P2V approximately 2 hours after SD offset and subsequently maintained a higher P2V value than the Pre phase at the same time point (Fig. 4C, Pre vs. RS, *** *P* = 3.1 × 10^-5^, two-way ANOVA). We calculated the maximum P2V values for each animal and compared these values between Pre and RS phases. However, we found no significant differences (Fig. 4D, Pre vs. RS, *P* = 0.49, bootstrap test for means). Furthermore, there were no significant differences in the SWS/REMS period or length of SWS or REMS bouts (Figs. 4E-G, SWS/REMS period, Pre vs. RS, *P* = 0.55, bootstrap test for means; SWS bout, Pre vs. RS, *P* = 0.86, bootstrap test for means; REMS bout, Pre vs. RS, *P* = 0.77, bootstrap test for means). These findings showed that, while the timing of elevated SWS/REMS regularity may be shifted by sleep needs, the underlying architectural characteristics persist.

**Figure 4.**
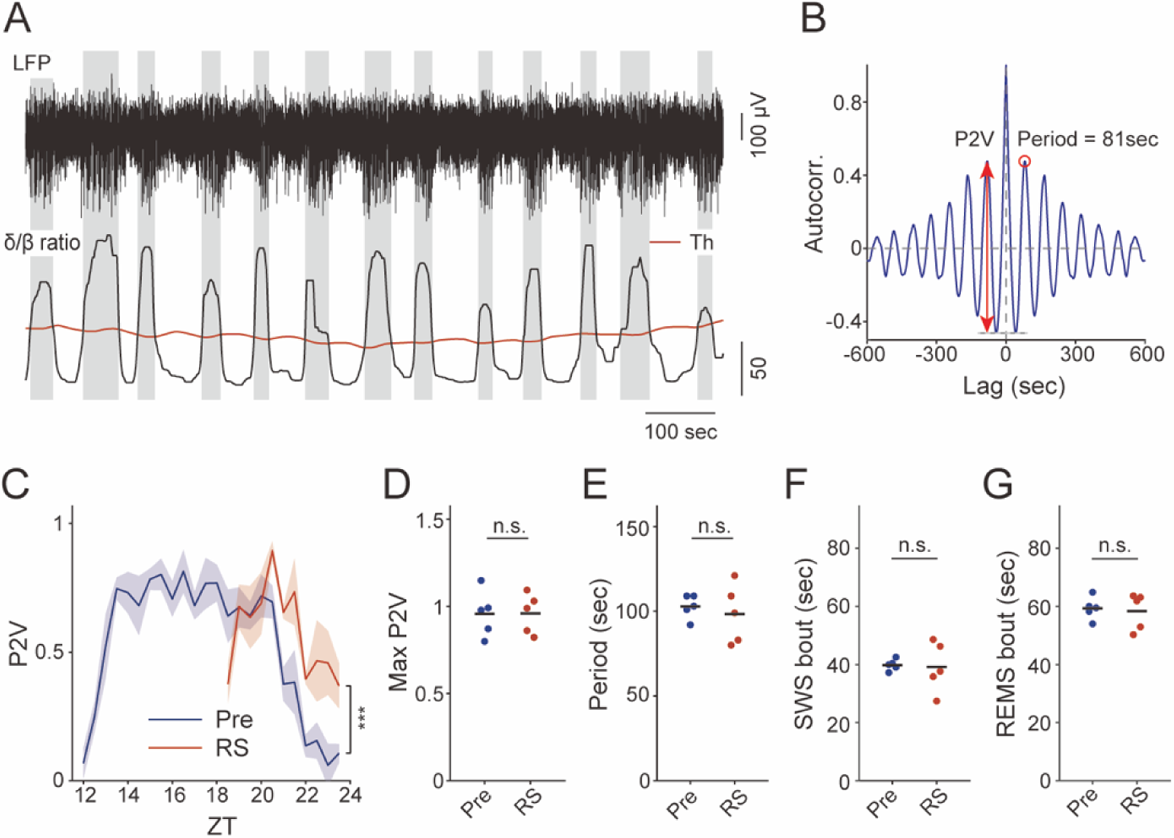
The regularity of REMS/SWS alternations is not affected by sleep deprivation. (A) LFP (top) and δ/β ratio dynamics (bottom) for 1000 sec during sleep. When the δ/β ratio is higher than the threshold (Th, red), readings are classified as SWS (gray) and otherwise REMS (white). (B) Representative autocorrelogram of the δ/β ratio dynamics. The δ/β dynamics shows a clear cycle with a period of 81 s (first peak, red circle). The difference between the first peak and the first valley (P2V) is marked by a red arrow. Differences are used as an index of the robustness of SWS/REMS alteration. (C) Transition of P2V value at night. N = 5 recordings each from 5 animals. Data are presented as mean ± SEM at 30 min intervals. (D) Max P2V values during the night. N = 5 recordings each from 5 animals. (E) SWS/REMS alternation period during which each animal’s P2V reaches its maximum value. N = 5 recordings each from 5 animals. (F) SWS bout length during which each animal’s P2V reaches its maximum value. N = 5 recordings each from 5 animals. (G) REMS bout length during which each animal’s P2V reaches its maximum value. N = 5 recordings each from 5 animals.

### Cortical lesions abolish rebounds in the LFP after SD

The reptilian cortex has a clear three-layered structure composed of diverse neuronal types, as seen in the mammalian cortex^20,21,22^. We next asked whether the three-layered reptilian cortex contributes to e-sleep homeostasis, as reported for six-layered neocortex in mammals^23^. To directly investigate the impact of the cortex on the rebound of sleep oscillations in the CLA/DVR, we bilaterally lesioned cortices, including the anterior dorsal cortex, the likely homolog of mammalian neocortex^22^ (Figs. 5A, B). During the baseline sleep, there were no significant differences between the intact and lesion groups in terms of the frequency and shape of ShWs, bout length of SWS and REMS, as well as regularity of transitions between the two sleep states (Figs. 5C-J, Amplitude/width of ShW, intact vs. lesion, *P* = 0.67, bootstrap test for means; ShW frequency, intact vs. lesion, *P* = 0.486, bootstrap test for means; max P2V, intact vs. lesion, *P* = 0.74, bootstrap test for means; period, intact vs. lesion, *P* = 0.45, bootstrap test for means; SWS bout, intact vs lesion, *P* = 0.25, bootstrap test for means; REMS bout, intact vs lesion, *P* = 0.10, bootstrap test for means). Following SD, however, throughout most of the RS, δ power rebound during SWS was suppressed in the lesion group, despite a slight rebound during the first 2 hours of RS (Figs. 5K, L, δ power ratio, intact vs. lesion, \*\*\**P* = 1.5 × 10^-6^, two-way ANOVA; δ power ratio in ZT 18.5-20, intact vs. lesion, *P* = 0.29, bootstrap test for means; δ power ratio in ZT 20-24, intact vs. lesion, \*\**P* = 1.1× 10^-3^, bootstrap test for means). This suppression relied on the decrease in the intensity of ShWs, rather than in the frequency (Figs. 5M-O, ShW amplitude ratio, intact vs. lesion, \*\*\**P* = 5.4 × 10^-7^, two-way ANOVA; ShW amplitude ratio in ZT 18.5-20, intact vs. lesion, *P* = 0.16, bootstrap test for means; ShW amplitude ratio in ZT 20-24, intact vs. lesion, \**P* = 0.036, bootstrap test for means; ShW frequency ratio, intact vs. lesion, *P* = 0.085, two-way ANOVA). In the lesion group, β power rebound during REMS also tended to be attenuated in RS (Fig. 5P, intact vs. lesion, \*\*\**P* = 1.1 × 10^-5^, two-way ANOVA). Furthermore, the δ rebound in the lesion group following the light-on phase was suppressed (Fig. 5Q, intact vs. lesion, \**P* = 0.01, bootstrap test for means). No significant difference was observed in β power after awakening (Fig. 5R, intact vs. lesion, *P* = 0.89, bootstrap test for means). These results suggest that the reptilian cortex is involved in sleep homeostasis, as in mammals, and regulates sleep-related neuronal activity in the CLA.

**Figure 5.**
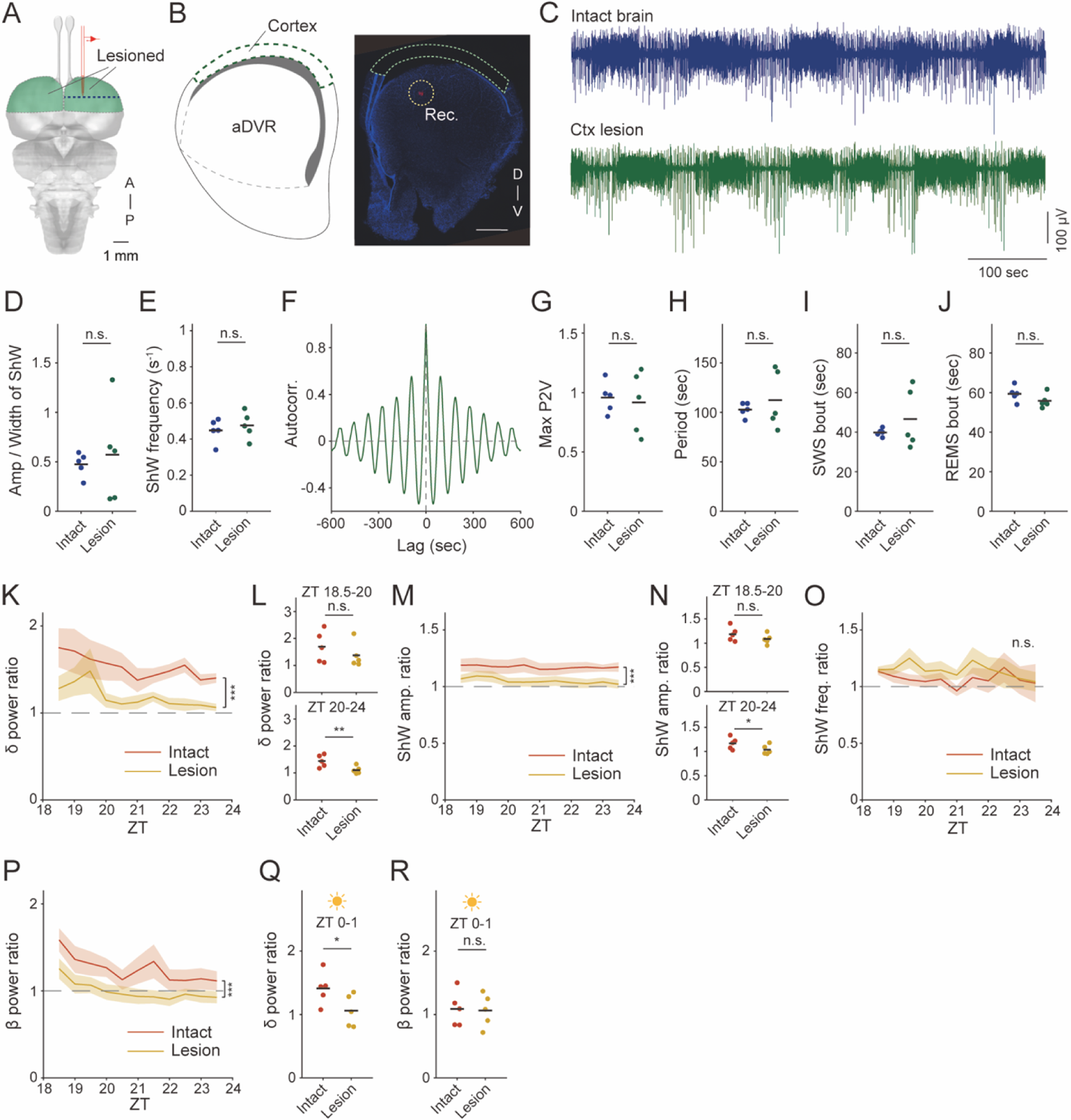
Bilateral cortical lesions attenuate the rebounds of LFP after SD (A) Schematic of the lesioned area (green). A, anterior; P, posterior. Coronal section outlined by blue dotted line is shown in B. (B) Coronal section with DAPI (blue) and DiI dye (red, white circle) to identify the recording site and cortical lesion. Scale bar, 1 mm. Rec, recording site; D, dorsal; V, ventral. (C) Representative LFP during sleep with the intact brain (blue) and cortex lesioned (green) (D) Averaged amplitude/width ratio of ShW during SWS. N = 5 recordings each from 5 animals. (E) ShW frequency in SWS. One hour during which each animal’s P2V reaches its maximum was quantified. N = 5 recordings each from 5 animals. (F) Autocorrelation from the cortex lesioned lizard, revealing a clear cycle. (G) Max P2V value in the night. N = 5 recordings each from 5 animals. (H) SWS/REMS alteration period in time showing the max P2V. N = 5 recordings each from 5 animals. (I) SWS bout length during which each animal’s P2V reaches its maximum value. N = 5 recordings each from 5 animals. (J) REMS bout length during which each animal’s P2V reaches its maximum value. N = 5 recordings each from 5 animals. (K) δ power ratio during SWS. N = 5 sessions from 5 animals each. Data are mean ± SEM at 30 min intervals. (L) δ power ratio at early phase (ZT 18.5–20) (top) and late phase (ZT 20–24) (bottom). N = 5 sessions from 5 animals each. (M) ShW amplitude ratio in SWS. N = 5 sessions from 5 animals each. The data are presented as mean ± SEM at 30 min intervals. (N) ShW amplitude ratio at early phase (ZT 18.5–20) (top) and late phase (ZT 20–24) (bottom). N = 5 sessions from 5 animals each. (O) ShW frequency ratio in SWS. N = 5 sessions from 5 animals each. Data are presented as mean ± SEM at 30 min intervals. (P) β power ratio in REMS. N = 5 sessions from 5 animals each. Data are presented as mean ± SEM at 30 min intervals. (Q) δ power ratio of the following morning (ZT 0–1). N = 5 sessions from 5 animals each (R) β power ratio of the following morning (ZT 0–1). N = 5 sessions from 5 animals each.

## Discussion

We have demonstrated electrophysiological signatures of sleep homeostasis in *Pogona,* revealing an increase in the sleep oscillations following sleep deprivation, suggesting that e-sleep homeostasis is a conserved feature among amniotes. Furthermore, we have shown that synaptic efficacy increases in the CLA/DVR after sleep deprivation, similar to what is known in mammals.

Our findings are partially inconsistent with a recent study on another lizard *Salvator merianae*, where SWS rebound was demonstrated by an increase in ShW frequency^17^. Our findings, along with the previous report, imply that the accumulation of sleep needs potentially affects the activity of neuronal populations responsible for low-frequency activity generation, regardless of variations in network mechanisms. It is also plausible that neuronal network in the forebrain relies on δ-band activity to alleviate sleep needs by modulating circuit excitability, including synaptic modification^24,25,26,27^.

In *Pogona*, sleep quantity was not affected by sleep deprivation. This is likely because the sleep-wake state in the lizards is strongly controlled by light input; light turning on during monophasic sleep (ZT 0) induces wakefulness. Considering the shift of peak timing of the P2V value during recovery sleep (Fig. 4C) and the increase in δ power observed the following morning during wakefulness, it is possible that extending sleep time might be observed if the light is kept turned off in the morning after sleep deprivation. Another possibility is that the sleep homeostasis in *Pogona* compensates for sleep deprivation by increasing sleep intensity rather than extending sleep duration. In some avian species, which are also diapsids like lizards, no increase in sleep quantity is observed during the recovery period, but there is an enhancement of slow waves^28, 29^. This suggests that differences in quantitative homeostatic responses to sleep deprivation might be species-specific.

Notably, cortical ablation did not significantly affect normal sleep but abolished the rebound after sleep deprivation. This suggests that the sleep need accumulates in the cortex even in the brains of reptiles, even though the sleep-related oscillations are not clearly observed in the cortical area. These data are consistent with recent findings highlighting the role of cortex in regulating the depth of SWS in mammals, including experiments that showed a diminished e-sleep rebound in mice, where synaptic transmission in a subset of cortical layer 5 neurons was inhibited^30,31,23^. Further investigations, including detailed observation of network dynamics in the cortex and CLA/DVR, as well as monitoring of synaptic plasticity during SD, will help us understand how the cortex integrates sleep need and regulates the activity of the CLA/DVR.

Finally, our study highlights the importance of conducting sleep research on non-model organisms. Quantifying the impact of cortical lesions on e-sleep homeostasis in mammals, where sleep stages are classified based on cortical activity, poses inherent challenges. However, the recent discovery of monophasic sleep in non-avian reptiles like *Pogona*, which possess a layered cortex and exhibit sleep stages characterized by neuronal activity in distinct regions outside the cortex, presents a unique opportunity to reveal the specific effects of the cortex on sleep homeostasis^5^. Given the remarkable diversity in the phenotypic characteristics of sleep across species, further investigations of sleep patterns in a wide range of animal species may help elucidate underlying common principles that govern sleep.

## Methods

### Animals

All procedures involving animal use complied with the guidelines of the National Institutes of Health and were approved by the Animal Care and Use Committee of the Hokkaido University (21-0092).

Lizards (*Pogona vitticeps*) of either sex, weighing 150–300 g, were obtained from breeders (Lab K, Sapporo, Japan). They were housed individually in reptile terraria (45×45×30 cm) and kept on a 12 h light/dark cycle with spotlights for basking (ZT 3-9). They had access to water *ad libitum* and were fed vegetables (bok choy, carrots, and lettuce) and proteins (live crickets) every other day. A powder supplement of calcium and vitamin D3 (EXOTERRA; GEX, Osaka, Japan) was added to the vegetables.

### Surgery

On the day before surgery, the lizards were given meloxicam (0.2 mg kg^−1^ *s.c.,* Boehringer Ingelheim animal health Japan, Tokyo, Japan) and habituated inside a sleep recording chamber (30×30×30 cm) within a soundproof box until the next morning. On the day of surgery, the lizards were placed in a box filled with isoflurane (Mylan Inc., PA, USA). After ensuring deep anesthesia, topical anesthesia (Sandoz Pharma, Basel, Switzerland) was sprayed around the trachea, and the tracheal cannula was inserted. Thereafter, the animals were placed on a stereotaxic instrument, and anesthesia was maintained with isoflurane (0.5-3 vol%). During surgery, body temperature was maintained around 32°C using a heating pad. Heart rate was monitored using an ultrasonic blood flow detector.

The skin covering the skull was disinfected with a 7.5% povidone-iodine solution (Meiji, Tokyo, Japan), after which the skin was removed, and a craniotomy was performed around the parietal eye (approximately 8×5 mm). The dura and arachnoid layers covering the forebrain were removed using fine forceps, and the pia around the area of electrode insertion was gently removed. The exposed skull was covered with a layer of ultraviolet (UV)-hardening glue (Transbond; 3M, MN, USA), and the bare ends of two insulated stainless-steel wires (0.005 inch, A-M SYSTEMS, WA, USA) were hooked to the subdura to serve as reference and ground.

For electrode insertion (either silicon probes or the same wire as the reference and ground), the electrode was mounted on a microdrive (homemade using a 3D printer) and secured to a stereotactic adaptor. The electrode was coated with the orange/red fluorescent membrane stain DiI (Fujifilm Wako, Osaka, Japan) for later histological assessment of the electrode position. The electrodes were slowly inserted into the forebrain about 1–1.5 mm from the brain surface, targeting the CLA/DVR. The brain surface was covered with duragel (Neurotech, Cambridge, UK). After connecting the ground and reference, the skull and microdrive were tightly secured using UV-hardening glue and dental cement (Sun Medical Company, Ltd, Shiga, Japan). EMG wires were implanted in the neck muscle.

After surgery, lizards were subcutaneously injected with meloxicam (0.2 mg kg^−1^ *s.c.*), released from the stereotaxic instrument, and left on a heating pad until full recovery from anesthesia.

### Cortical lesion

After a craniotomy in the process of surgery, a part of the forebrain is exposed. The cortex has a sheet-like structure and located on the surface of the brain over the DVR. The bilateral cortices were surgically removed by micro-scissors without damaging any other brain regions including the DVR. Lesioned areas correspond to the likely homolog of neocortex and a part of hippocampus in mammals^22^. After cortical lesion, the electrode is directly implanted into the CLA/DVR so that its position is close to the control group animals. The subsequent process is the same as in the control animals.

### Recording

Following recovery, each lizard was placed inside a soundproof chamber (30×30×30 cm). The animal was left overnight to record its natural sleep and returned to its home terrarium 1 hour after the lights were turned on. The recording started at ZT 11 and stopped at ZT 1. Because the animal had access to water but not food within the chamber, it was fed after returning to its terrarium. Light and temperature conditions in the recording room were identical to those in the home terrarium.

Recordings were acquired using an acquisition board (Open Ephys, Lisbon, Portugal) and 32 Ch headstages (Intan technologies, CA, USA) as previously described^32^. Recordings were grounded and referenced against a reference wire. Signals were sampled at 1 kHz with a 0.1–100 Hz band-pass filter.

### Sleep deprivation

After obtaining baseline recordings, the lizard underwent sleep deprivation from ZT 11 to ZT18 by gentle handling and visual distraction (Supplementary Movie 1). When the lizards showed obvious signs of sleepiness, they were stimulated using a brush. As the frequency of stimulation had to be increased, crickets in the beaker were introduced to capture the lizard’s attention and keep it awake. The non-sleep deprivation group (NSD) was the control group, and LFP was recorded for two consecutive nights under standardized conditions.

### Histology

Lizards received deep anesthesia using isoflurane and ketamine (60 mg kg^−1^ *i.m.*, Fujita Pharm, Tokyo, Japan) until the corneal reflex was lost. After confirming the loss of corneal reflex, animals were perfused transcardially with more than 150 ml of cold phosphate-buffered saline (PBS) followed by 150 ml of 4% paraformaldehyde (PFA). After decapitation, the brain was carefully removed, and the dura and pia on its surface were peeled off using forceps. The brain sample was fixed with 4% PFA overnight at 4°C and subsequently immersed in 30% sucrose at 4°C. 100 μm-thick coronal brain sections were made using a sliding microtome (HM450; Epredia, NH, USA) at −21°C. Sections were immersed in mounting medium with DAPI (H1500; Vector Laboratories, CA, USA). Sections were photographed using a BZ-X810 microscope with a 2× objective lens (Keyence, Osaka, Japan). Images were processed with BZ-II analysis application and ImageJ (NIH, MD, USA).

### Sleep stages

The data underwent fast Fourier transform (FFT) analysis with a 10 sec window and 1 sec step from 0.1–30 Hz. The δ/β ratio was calculated every 1 sec bin by dividing the average spectrum over frequencies between 0.1 and 3 Hz by the average spectrum over frequencies between 10 and 30 Hz. The δ/β ratio was smoothed with a 20-sec median filter then a 601-sec moving average. For the night data, the median filtered δ/β ratio was compared with the moving average δ/β ratio for each bin to determine the sleep stage; cases wherein the median filtered δ/β ratio was higher than the moving average δ/β ratio were designated as SWS and the others as REMS. For EMG recording, the electrostatic noise was removed by visually setting a threshold. Then, the root mean square (RMS) was calculated with a 60-sec window. Electrostatic noise was removed by setting a visual threshold on the LFP plot.

### Autocorrelation

SWS/REMS alteration period was calculated using the autocorrelation of the δ/β ratio. The raw δ/β ratio was smoothed with a 3601-sec moving average. The time point with the maximum value in the night was defined as δ/β peak time. The autocorrelation was calculated for a 4-hour window centered on the δ/β peak time. From the autocorrelation function, the global oscillation period and antiphase lag were extracted by locating the first positive and negative nonzero peaks, respectively. A scrolling autocorrelation was calculated on the δ/β time series (1800-sec bins). For each bin, the local period and anti-phase lag were calculated by taking the maximum and minimum readings during a 20-sec interval surrounding the global period and global anti-phase lag, respectively. Peak-to-valley (P2V) was defined as the difference between the first peak and first valley of the autocorrelogram. The local period, wherein each animal’s P2V reaches its maximum value, was defined as the SWS/REMS cycle period.

### Sharp-wave detection

Data recorded during SWS was band-pass-filtered between 0.5–4 Hz. The −2 standard deviation (s.d.) of all data in SWS was set as the threshold for detecting ShW. Troughs below this threshold were identified as ShWs. ShW amplitude was defined as the absolute value of the ShW trough. ShW frequency was calculated as the number of ShWs per second of SWS.

### *In vitro* electrophysiology

Adult lizards were deeply anaesthetized with isoflurane, ketamine (60 mg kg^−1^). After loss of the corneal reflex, the animals were decapitated, and the brain was coronally sliced (600 μm thick) in an ice-cold oxygenated artificial cerebrospinal fluid (aCSF) consisting of (in mM) 27 NaCl, 26 NaHCO_3_, 1.6 KCl, 1.24 KH_2_PO_4_, 1.3 MgSO_4_, 2.4 CaCl_2_, and 10 D-glucose bubbled with carbogen gas (95% O2, 5% CO2) using a vibratome (VT1200, Leica Biosystems, Wetzlar, Germany). Slices were allowed to recover for at least 30 min while submerged in a chamber filled with oxygenated aCSF at room temperature. Long-shank patch pipettes (4–6 MΩ) were pulled from borosilicate glass with a Sutter P1000 electrode puller. Pipettes were filled with internal solution (127 mM CsMeSO_4_, 8 mM CsCl, 10 mM HEPES, 1 mM MgCl_2_, 10 mM phosphocreatine-Na_2_, 4 mM MgATP, 0.3 mM NaGTP, and 0.2 mM EGTA (pH 7.2–7.3, 280–295 mOsm).

Experiments were carried out on an upright Olympus BX51WI microscope with 10× and 40× water-immersion objectives and cells were patched under visual guidance. mEPSCs were recorded at a holding potential of −70 mV in the presence of tetrodotoxin (1 μM, Alomone labs, Jerusalem, Israel). mEPSCs were detected using an in-house MATLAB program and were defined as inward currents with amplitudes > 2.5 s.d. The series resistance was monitored, and if it exceeded 30 MΩ, the data were discarded. Data were sampled at 20 kHz and filtered at 2 kHz using a Double IPA amplifier (Sutter Instruments, CA, USA)

### Statistics

All analyses were done using MATLAB (R2022a; MathWorks, MA, USA). Statistical details of the experiments are presented in the text. For the time series comparison of the groups, a two-way ANOVA was performed with the graph representing the mean (line) ± standard error of the mean (SEM, shade). To compare the groups, a bootstrap test for means was performed^33^. In the bootstrap test, data in each group were re-sampled for 1,000,000 times with duplicates allowed, and the means of each simulated data were calculated by “bootstrap” function in MATLAB (=1,000,000 means in each group). Then, the differences between the means of each group were calculated, and the number of times these differences were greater than zero was counted. Based on this number, a p-value was calculated.

In the graph, the dots represent individual values, and the black line represents the mean of each group. In all comparisons, a p-value of < 0.05 was considered statistically significant. n.s.; not significant.

## Acknowledgements

We thank Drs. Lorenz A. Fenk and Gilles Laurent for discussion, comment on the manuscript, and confirmation of data reproducibility; all the members of the Norimoto Laboratory for their daily discussions and advice. This work was funded by JSPS KAKENHI (JP22K15369) and grant form Hirose Foundation to Z.Z., Grant-in-Aid for JSPS Fellows (JP23KJ0055) to S.H., KAKENHI (JP23K14225 and JP22K20675), the Akiyama Life Science Foundation and the Sasakawa Scientific Research Grant to S.T.Y., KAKENHI (JP23KJ0004, JP24K18155) to R.K., and JST PRESTO (JPMJPR2048), AMED (22wm0525003s0202), KAKENHI (JP23K18251, JP24K02058, JP 23K18251, JP24H01996), the Murata Science Foundation, the Uehara Memorial Foundation, the Mochida Memorial Foundation for Medical and Pharmaceutical Research, a Grant for Basic Science Research Projects from the Sumitomo Foundation, the Astellas Foundation for Research on Metabolic Disorders, the Nakajima foundation, the Naito foundation, and the Toray Science and Technology Grant to H.N.

## Author Contributions

S.H., S.T.Y., and H.N. designed the study. S.H., S.T.Y., and R.K. performed the experiments. S.H., S.T.Y., and R.K. analyzed the data and prepared the figures. F.M. assisted with the sleep deprivation experiment. Z.Z. assisted with the analysis. All authors discussed and interpreted the results. S.H., Z.Z., and H.N. wrote the manuscript with input from all authors.

## Data and code availability

Datasets are available from the corresponding author upon reasonable request.

