## Supplementary Figure1 for "Sleep homeostasis in lizards and the role of cortex"

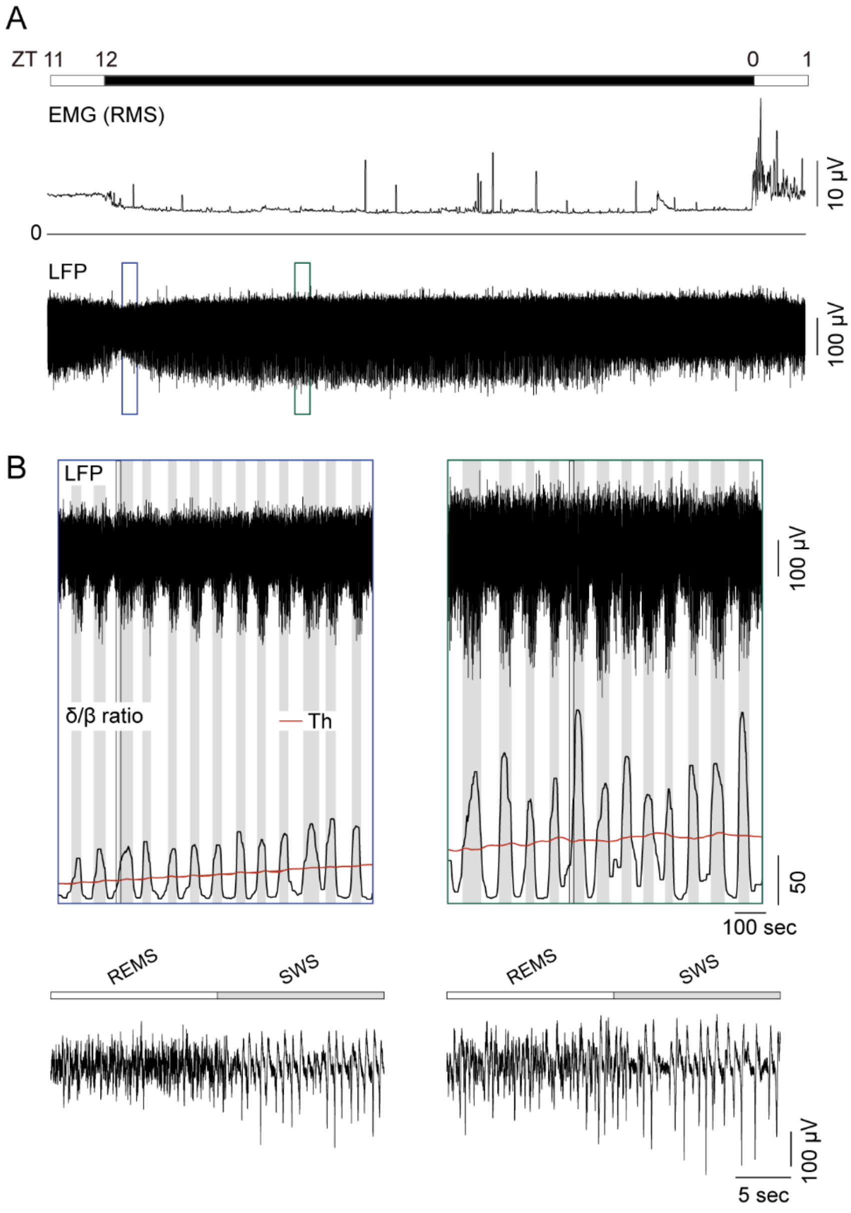


**Figure S1. Electrophysiological sleep in a night and sleep stage analysis**

(A) Representative EMG (RMS) and LFP in a recording period. Note that EMG is toned down during dark periods with muscle twitching also observed. The enclosed areas in LFP are magnified in B.

(B) Top: LFP magnified from A and δ/β ratio. The red line represents the threshold (Th) for classifying the sleep stage; when the δ/β ratio exceeds the red line, it is classified as SWS (gray), and otherwise REMS (white). The enclosed areas are magnified in the bottom. Bottom: SWS/REMS alteration point surrounded in the top.
